# Theory of Temporal Pattern Learning in Echo State Networks

**DOI:** 10.1101/2025.06.23.661158

**Authors:** Vincent Hakim, Alain Karma

**Affiliations:** Laboratoire de Physique de l’Ecole Normale Supérieure, CNRS, Ecole Normale Supérieure, PSL University, Sorbonne Université, Université Paris-Cité, Paris, France; Physics Department and Center for Interdisciplinary Research in Complex Systems, Northeastern University, Boston, MA, USA

## Abstract

Echo state networks are well-known for their ability to learn temporal patterns through simple feedback to a large recurrent network with random connections. However, the learning process itself remains poorly understood. We develop a quantitative theory that explains learning in a regime where the network dynamics is stable and the feedback is weak. We show that the dynamics is governed by a finite number of master modes whose nonlinear interactions can be described by a normal form. This formulation provides a simple picture of learning as a Fourier decomposition of the target pattern with amplitudes determined by nonlinear interactions that, remarkably, become independent of the network randomness in the limit of large network size. We further show that the description extends to moderate feedback and recurrent networks with multiple unstable modes.

---

Twenty years ago, Jaeger and Haas [1] presented a remarkable way of mimicking an unknown nonlinear dynamical system that output a given time series. They showed that feeding back into the network an appropriate linear combination of its unit activities can drive it to produce any temporal pattern. The development of these so-called Echo-State networks (ESNs), has given rise to a very active engineering field [2, 3] as well as much further work in neuroscience. In this latter context, it was shown that learning the suitable feedback could be produced “online” by comparing at each time the output of the network to the signal [4]. This has led to a flourishing activity in which training dynamical networks helps to understand the dynamics of real biological neural networks[5]. However, the principles that underlie the success of ESNs at learning a variety of temporal patterns have remained incompletely understood, in spite of several previous attempts e.g. [6–9]. Here, we analyze the prototypical ESN of N recurrent units described by

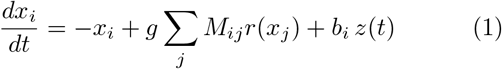

where *x*_*i*_ quantifies the activity of the i-th unit, **M** is a Gaussian random matrix with non-diagonal elements of variance 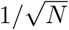 and with zero diagonal elements corresponding to no self-interactions. The nonlinear “rate” function *r*(*x*) measures the output of each unit, taken here, as usual, to be tanh(*x*) for simplicity. The linear combination of outputs *z*(*t*) = ∑_*j*_ *w*_*j*_*r*[*x*_*j*_(*t*)]) serves both as a readout of the network activity and as a feedback on the network dynamics through the fixed random vector **b** with elements *b*_*i*_ of order one. Learning consists of finding the weights *w*_*j*_ such that a desired periodic function *f* (*t*) of period *T* is reproduced, as best as possible, by the readout *z*(*t*), autonomously produced by the network dynamics of Eq. (1). One usual way of computing the weights *w*_*j*_ is to minimize the loss function

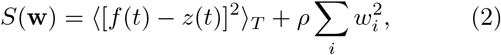

where the brackets denote time averaging over one period, 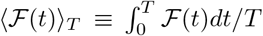, and the second term on the right-hand-side (r.h.s.) is a L2 regularization that controls the magnitude of the weight vector **w** [1]. The weights corresponding to the unique minimum of *S*(**w**) satisfy the linear equation

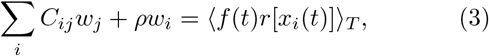

where **C** is the correlation matrix between the rates of the network units, namely *C*_*ij*_ = ⟨*r*[*x*_*i*_(*t*)]*r*[*x*_*j*_(*t*)] ⟩ _*T*_ .

While different approaches have been developed to compute **w** [1, 4], we use here the simplest one [1]. The correlation matrix and the r.h.s. of Eq. (3) are evaluated by using, instead of the autonomous dynamics **x**(*t*), the forced dynamics 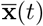 obtained by replacing *z*(*t*) by *f* (*t*) in Eq. (1), implicitly assuming that learning is successful (*z*(*t*) = *f* (*t*)). We then run the autonomous dynamics of Eq. (1) with *z*(*t*) = ∑_*j*_ *w*_*j*_*r*[*x*_*j*_(*t*)]) as a self-consistency check of this assumption. This assumption turns out to be satisfied for all present computations that use a small regularization (*ρ* = 10^−10^) ensuring that the cost function is minimized for *z*(*t*) very close to *f* (*t*). Importantly for what follows, even when *z*(*t*) ≈ *f* (*t*) over one period, the learned nonlinear limit cycle of the autonomous dynamics can itself be linearly stable or unstable with *z*(*t*) remaining close to or deviating from *f* (*t*), respectively, over several periods.

A classic result of random matrix theory states that the eigenvalues of large matrices **M** are uniformly distributed in the unit disk [10]. To understand the learning process, we focus first on a regime where the dynamics without feedback is linearly stable (*g <* 1) and the fee back is weak (*f* (*t*) ≪ 1). This regime is made analy cally tractable by the fact that the learned dynamics slaved to a few small amplitude “master modes”, which weakly interact nonlinearly to produce stable or unstable limit cycles. As a result, the dynamics of Eq. (1) can reduced to the evolution equations for the complex a plitudes of those modes using a normal form approach dating back to Poincaré and widely used to reduce the dimensionality of nonlinear dynamical systems [11–1 here from *N* ≫ 1 to a small number of modes.

To illustrate the existence of these modes, we first co pare in Fig. 1 the results of learning a truncated Fourier sine series of a sawtooth function *f* (*t*) = *A*[sin(*ωt*) − sin(3*ωt*)*/*9 + sin(5*ωt*)*/*25] (*A* = 0.1 and *ω* = *π/*10) with both the nonlinear dynamics of interest, *r*(*x*) = tanh(*x* and its linear approximation, *r*(*x*) = *x*; these two choices produce different **w** due to the presence and absence of nonlinearity through *r*(*x*) in Eqs. (1) and (3), respectively. However, remarkably, they yield nearly indistinguishable autonomous network dynamics (Eq. (1) with *z*(*t*) = ∑_*j*_ *w*_*j*_*r*[*x*_*j*_(*t*)])) reproducing the target sawtooth (Fig. 1a). They also yield very similar stability spectra of the autonomous dynamics of Eq. (1) linearized around **x** = 0, corresponding to *d***x***/dt* = **Lx** where

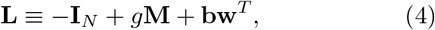

and **I**_*N*_ denotes the N-dimensional identity matrix. As shown in Fig. 1b, the spectrum of the matrix **L** with **w** learned with *r*(*x*) = *x* displays three pairs of complex conjugate eigenvalues with vanishing real parts and imaginary parts exactly equal to the frequencies *ω*_*n*_ = ±*nω*, (*n* = 1, 3, 5) of the sawtooth. In this case, the autonomous dynamics is purely linear and any superposition of these six modes is a solution of the autonomous dynamics consistent with the findings of previous studies [16–18]. The sawtooth function is obtained by precisely choosing the amplitudes of these six modes to produce the desired output superposition, *z*(*t*) = ∑_*n*_ *Z*_*n*_ exp(*iω*_*n*_*t*), where the *Z*_*n*_ match the amplitudes *f*_*n*_ of the target function *f* (*t*) = ∑_*n*_ *Z*_*n*_ exp(*iω*_*n*_*t*) as detailed in the *End Matter* section. However, as a result of linearity (*r*(*x*) = *x*), the autonomous dynamics is marginally stable, i.e. any perturbation changes the amplitudes *Z*_*n*_ and thus changes the output function *z*(*t*). In contrast, with nonlinearity (*r*(*x*) = tanh(*x*)), one or more pairs of complex conjugate eigenvalues have a small positive real part and *Z*_*n*_ ≈ *f*_*n*_ must emerge as a fixed point of the slow dynamics governing the evolution of the complex amplitudes *Z*_*n*_(*t*) over a time scale larger than *T* . Understanding learning is now reduced to answering the two basic questions: How do fixed points emerge? What controls their stability?

**FIG. 1.**
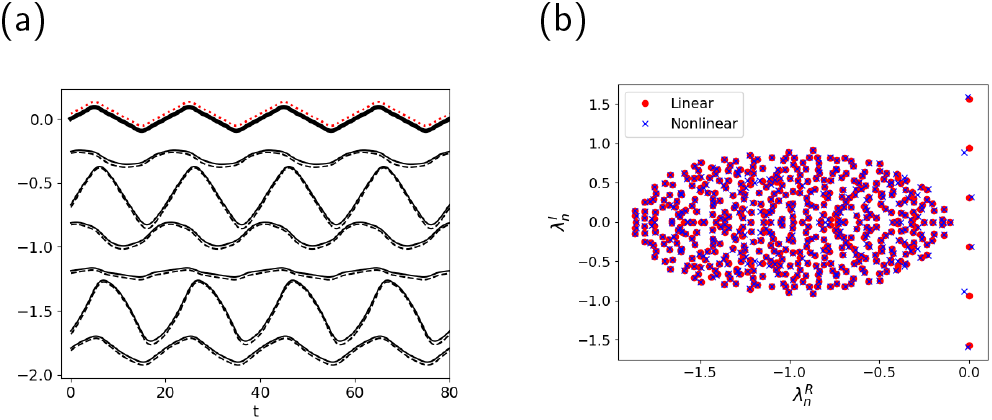
An example of ESN learning in the stable (*g* = 0.9) network regime. (a) The autonomous dynamics of the network (thick solid dark line) reproduces after learning the target “sawtooth” (red dotted line slightly shifted upward for visibility). Examples of network unit activities are also shown both for the nonlinear (thin solid dark lines) and linear (dashed dark lines slightly shifted upward for visibility) cases. (b) Spectrum of the associated linear dynamics (*N* = 500) both in the nonlinear (red circles) and linear (blue crosses) cases. Note the few separated eigenvalues with zero (linear case) or small (nonlinear case) real parts, which correspond to the frequencies of the sawtooth. Eq. (3) for both forced and autonomous dynamics is solved using a fourth-order Runge-Kutta algorithm with *dt* = 0.1.

The simplest possible setting to address these questions is provided by the case of a single frequency, *f* (*t*) = *f*_1_ exp(*iωt*) + *c*.*c*., which is illustrated in Fig. 2a for a pure cosine 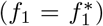 of amplitude 2*f*_1_ [19]. The linear operator **L** after learning (Fig. 2b,), has two slow modes with complex conjugate eigenvalues equal to *λ*_1_ and 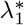, close to ±*iω*. The network activity at the linear level is taken to be **x**(*t*) = *A*_1_(*t*) exp(*iωt*) **u**_1_ + *c*.*c*. where *A*_1_ is arbitrary and **u**_1_ is the right eigenvector of **L** corresponding to the eigenvalue *λ*_1_. At the next nonlinear order, the slow evolution of *A*_1_(*t*) is determined by the classical Poincaré-Hopf normal form. It is obtained from the non-appearance of resonant terms in the dynamics [11–14], which yields the amplitude equation

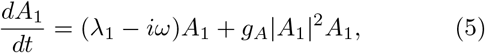

where *A*_1_ evolves slowly on the time scale of the period *T* since |*λ*_1_ − i*ω*| ≪ 1 when |*f*_1_| ≪ 1. The form of Eq. (5) is determined by the symmetries of the full network dynamics (Eq. (1)). Namely, Eq. (5) is the lowest order equation invariant under time translation, *A*_1_ → exp(*iϕ*)*A*_1_ and under the reflection **x** → −**x**, which translates into *A*_1_ → −*A*_1_. The value of the constant *g*_*A*_ is obtained as,

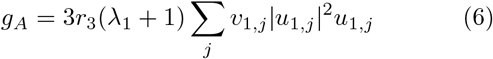

where **u**_1_ and **v**_1_ are right and left eigenvectors of **L** for the eigenvalue *λ*_1_, with the normalization 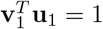, and *r*_3_ = −1*/*3 is the coefficient of the cubic nonlinearity of *r*(*x*) = tanh *x* = *x* + *r*_3_*x*^3^ +… as detailed in *End Matter*. The evolution of *z*(*t*) = *Z*_1_(*t*) exp(*iωt*)+*c*.*c*, is obtained in turn by projecting the network activity with the readout vector **w**,

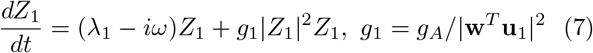

**FIG. 2.**
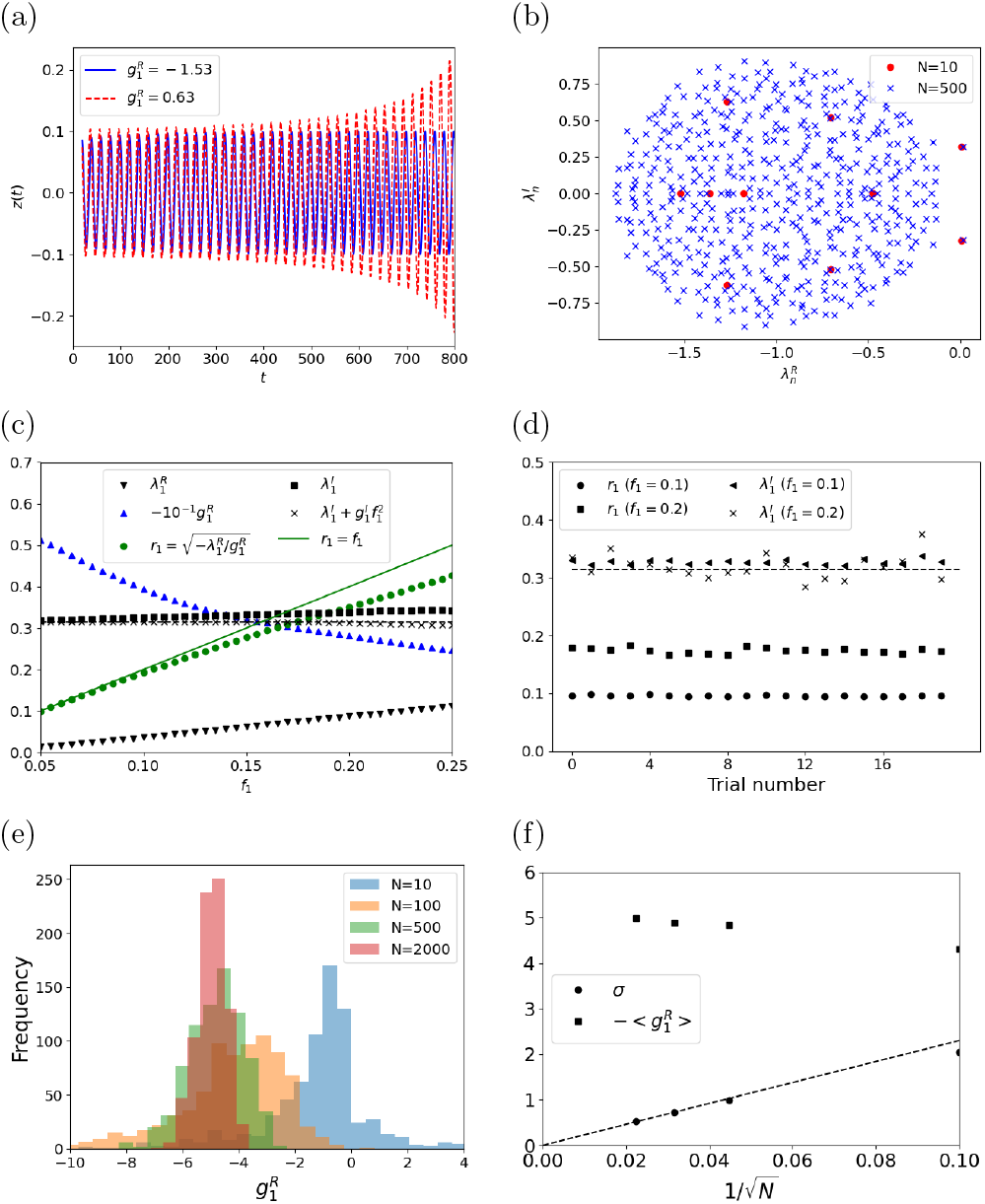
Learning a single cosine function 2*f*_1_ cos(*ωt*) (*ω* = *π/*10).(a) Two examples of autonomous dynamics after learning (*f*_1_ = 0.05) which can be stable (blue solid line) or unstable (red dashed line). The real part of 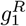 (Eq. (7)) is indicated in each case. (b) Examples of **L** spectra after learning fro a small (*N* = 10) and a large network (*N* = 500). (c) For a given matrix **M**, it is displayed after learning, as a function of the amplitude 2*f*_1_, the real and imaginary parts of the eigenvalue *λ*_1_ and the coefficient *g*_1_ (Eq. (7), as well as the predicted amplitude of the autonomous limit cycle *r*_1_ as well its nonlinearly corrected frequency (Eq. (8)). The amplitude of the read-out *z*(*t*) after learning is indistinguishable from *f*_1_ (solid green line). (d) Predicted amplitude *r*_1_ and non-corrected frequency 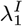 of the autonomous dynamics after learning for a set of 20 matrices **M**, for two amplitudes *f*_1_.(e) Histogram of the values of the real part 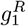 for different network sizes. (f) Mean value 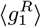 and standard deviation *σ* of the histogram in (e) as a function of network sizes.

The real and imaginary part of the linear dynamics eigenvalue, 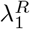 and 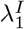 and of the Poincaré-Hopf coefficient *g*_1_, i.e 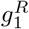 and 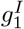 determine whether Eq. (7) possesses a limit cycle and what its amplitude *r*_1_ and frequency *δω*_1_ are. Namely when 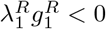, these are

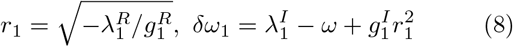

For learning the correct fixed point, independent of its stability discussed below, the choice of **w** which controls both *λ*_1_ and *g*_1_, should be such that the output *z*(*t*) exactly matches the target pure cosine 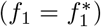 of amplitude 2*f*_1_ and frequency *ω*, which implies the two equalities *r*_1_ = *f*_1_ and *δω*_1_ = 0.

To obtain the coefficient *g*_1_ and test these results, we computed the left and right eigenvectors, **u**_1_ and **v**_1_, of the matrix **L** with **w** solution of Eq. (3). The results are illustrated in Fig. 2c for a given matrix **M** and cosine functions of different amplitudes 2*f*_1_. The slow mode eigenvalue *λ*_1_ starts from ±*iω* for a vanishing *f*_1_ and has a growing real part as *f*_1_ increases. The corresponding coefficient *g*_1_ also varies with *f*_1_ and combines with 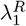 such that the resulting amplitude *r*_1_ of the autonomous limit cycle matches *f*_1_, in accordance with Eq. (8). This is also true for the frequency of the limit cycle. As shown in Fig. 2c, the nonlinear departure of the frequency from *ω* due to the imaginary part 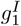 of *g*_1_ is compensated by the departure of the imaginary part of 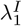 from *ω* so that *δω*_1_ ≃ 0. The close agreement between the predicted amplitude *r*_1_ of the limit cycle and the desired amplitude *f*_1_ is observed independently of the chosen matrix **M** as illustrated in Fig. 2d.

The above conditions on *r*_1_ and *δω*_1_ ensure that the proper feedback has been learned, such that the autonomous dynamics display a limit cycle of the right amplitude and frequency. However, they do not ensure the stability of this limit cycle and full learning success. In this one-frequency case, the existence of a limit cycle (Eq. (8)), requires the real part *λ*_*R*_ of the slow mode and *g*_1_ to be of opposite signs. Stability of the limit cycle simply corresponds to 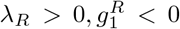 whereas for 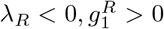 any small perturbation of the amplitude *A*_1_ makes it grow away from *f*_1_. Learning can in principle produce both cases. As illustrated in Fig. 2a for networks of size *N* = 10, starting from randomly drawn matrices **M**, autonomous dynamics with unstable limit cycle are often produced. However, for networks sizes of 100 or higher, quite remarkably, instability is almost never observed. In order to better understand, this surprising growing success of learning with the network size, we computed the distribution of Poincaré-Hopf coefficients *g*_1_ given by Eq. (7) for ensemble of random matrices of different sizes *N* . The corresponding histograms of *g*_1_ are shown in Fig. 2e. The histogram clearly shrinks around their negative mean value as *N* grows. As quantified <u>in</u> Fig. 2f, the standard deviation of *g*_1_ decreases as 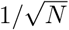 around a mean value that extrapolates for *N* → + ∞ to the negative value 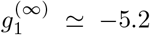. This clearly explains why positive Poincaré-Hopf coefficients are less and less frequently found as *N* grows. Although, the results of Fig. 2e,f are shown for *f*_1_ = 0.05, we generally observed this behavior for all amplitudes in the range 0 ≤ *f*_1_ ≤ 0.25. It is worth noting that this remarkable stability of the learned dynamics for large networks which holds independently of the considered random network, depends on the saturating character of the tanh cubic nonlinearity, *r*_3_ *<* 0, as shown by Eq. (6). For a rate activation with *r*_3_ *>* 0 on the contrary almost no limit cycle would be stable for large network sizes.

The theory is easily applied to describe the learning of more complex functions with a finite number of frequencies as the sawtooth of Fig. 1. We treat here the case of a general two-frequency function *f* (*t*) = [*f*_1_ exp(*iωt*) + *f*_2_ exp(*i*2*ωt*) + *c*.*c*] with the case 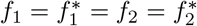 illustrated in Fig. 3. In this case, learning with a linear rate function with a vanishingly small regularisation produces an operator **L** with 4 slow modes with imaginary parts equal to ±*iω* and ±*i*2*ω*. Learning from the full Eq. (3), displaces the two couples of eigenvalues to 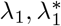 and 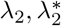, as shown in Fig. 3a. The network output function is described as *z*(*t*) = [*Z*_1_ exp(*iωt*) + *Z*_2_ exp(*i*2*ωt*) + *c*.*c*] with *Z*_1_ and *Z*_2_ “slow” time-dependent functions, the dynamics of which depends both on the displaced linear eigenvalues and the nonlinear interactions between the slow modes. Their evolution is governed by the following normal form,

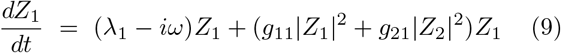

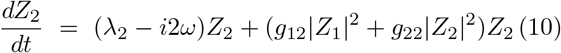

with the constants *g*_*ij*_ given by expressions similar to Eq. (7) for *g*_1_, as detailed in *End Matter*. As in the previous example (Eq. (7)), the cubic lowest nonlinear terms that appear are restricted by symmetries, namely time translation invariance, *Z*_1_, *Z*_2_ → exp(*iϕ*)*Z*_1_, exp(*i*2*ϕ*)*Z*_2_, and the inversion, *Z*_1_, *Z*_2_ → −*Z*_1_, −*Z*_2_ coming from the use of tanh activation in Eq. (1).

**FIG. 3.**
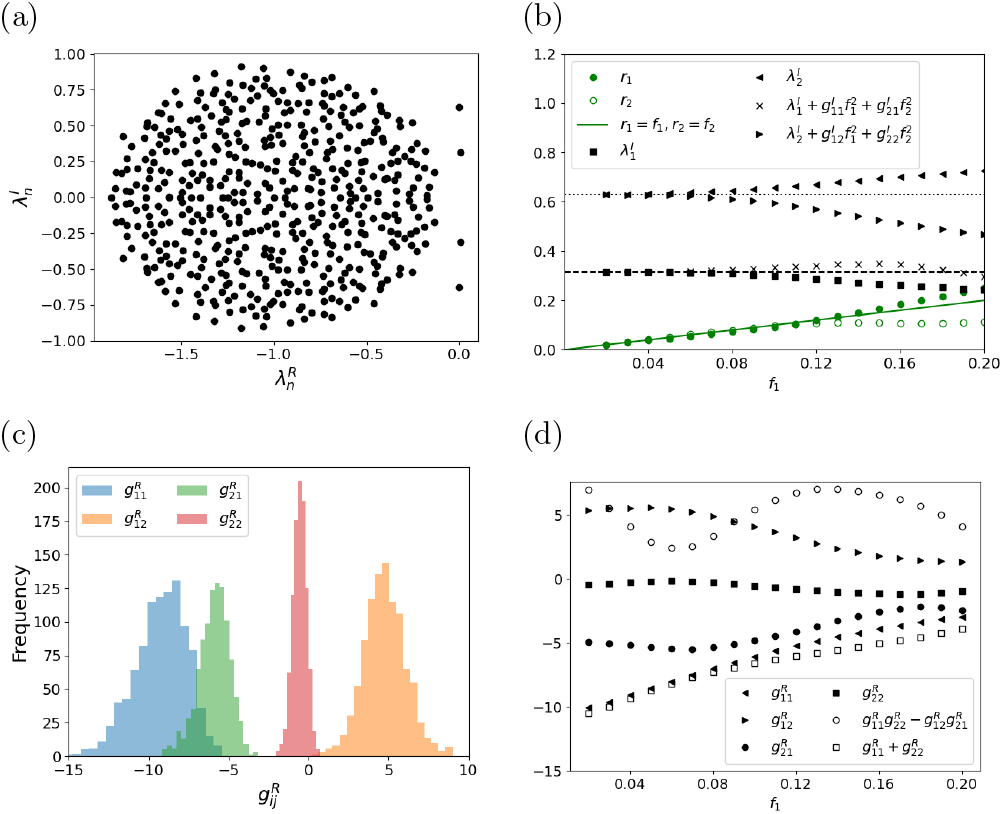
Learning a two-frequency function *f* (*t*) = 2*f*_1_[cos(*ωt*) + cos(2*ωt*)] with *ω* = *π/*10. (a) Example of **L**-spectrum (Eq. (4)) after learning (*f*_1_ = 0.025, *N* = 500). (b) Results of learning for different amplitudes *f*_1_ for the same matrix **M** as in (a). The amplitudes *r*_1_ and *r*_2_ as obtained from Eq. (11) are shown as well as the imaginary parts 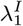 and 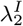 of the two dominant eigenvalues, together with the corresponding nonlinearly corrected imaginary parts (Eq. (12)). The target frequencies *ω* (dashed line) and 2*ω* (dotted line) are also shown. (c) Histograms of the real parts of the coefficients *g*_*ij*_ (Eq. (22)) obtained for learning with *f*_1_ = 0.025, *N* = 500, and 200 different matrices **M**. (d) The real parts of the coefficients *g*_*ij*_ are shown for different amplitudes *f*_1_ together with the limit-cycle stability conditions (Eq. (13)) (same simulations as in (b)). Note that the stability conditions (Eq. (13)) are always satisfied.

A limit cycle with amplitudes *r*_1_ = |*Z*_1_| and *r*_2_ = |*Z*_2_| exists when the following conditions are satisfied,

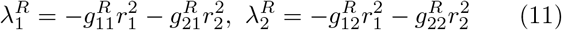

In order to reproduce the target function *f* (*t*), the solutions *r*_1_ and *r*_2_ of Eq. (11) should obey *r*_1_ = |*f*_1_|, *r*_2_ = |*f*_2_| . In order to obtain the correct frequencies of the two modes, the imaginary parts of linear modes 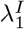 and 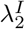 should moreover compensate the nonlinear frequency corrections,

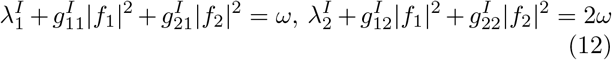

For large *N* (i.e. *N* = 500), learning is almost always successful and both the amplitude and phase conditions are found to be well-obeyed, as illustrated in Fig. 3b for a given matrix **M** and varying amplitude. As in the simple case of the cosine function (Eq. (7)), Eq. (11) and (12) are not sufficient to ensure the stability of the autonomous dynamics limit cycle. Taking *Z*_1_ = *r*_1_ exp(*iϕ*_1_) and *Z*_2_ =, *r*_2_ exp(*iϕ*_2_), a simple analysis shows that the fixed point amplitudes *r*_1_ = *f*_1_ and *r*_2_ = *f*_2_ are linearly stable when the two following conditions are obeyed

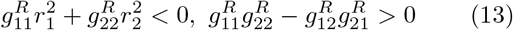

Quite remarkably for *N* = 500 or larger, the limit cycle is found to be always stable [20] and these relations always satisfied as shown in Fig. 3c for an ensemble of matrices **M** for the same two-cosine function *f* (*t*), and in Fig. 3d for the same matrix **M** and varying the amplitude of this function. As in the simpler single frequency case, this results from the *g*_*ij*_ coefficients tending to well-defined limiting values in the large network size limit, independently of the random network interaction matrix **M**. In particular, the limiting values of 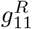 and 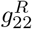 are negative while 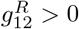 and 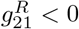 are of opposite signs, ensuring the two conditions (13) for any values of the limit cycle amplitudes *f*_1_ and *f*_2_. Of note, these limiting values ensure that 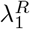 is positive and can drive the growth of the limit cycle. While this asymptotic stability for the one- and two-mode functions appears quite remarkable, preliminary calculations suggest that stability for functions with more modes depends on the specific choice of function and is not as robust.

We have focused above on the case when the recurrent without feedback is stable (*g <* 1). However, the normal form description can also be applied in the case when *g >* 1 and the dynamics without feedback has a number of unstable modes. Fig. 4a shows the whole spectrum of the matrix **L** when successfully learning a simple cosine function with *g* = 1.1. It comprises a pair of unstable modes with imaginary parts close to 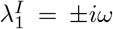 and a positive real part 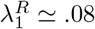 similarly to spectra obtained for *g <* 1 (Fig. 2). However, for *g* = 1.1, there are seven other linearly unstable modes as seen in the spectrum magnification (Fig. 4b). Contrary to naive expectations these modes do not prevent the stable learning of the sine function. At the weakly nonlinear level, the normal form approach developed here provides the (lowest-order) interaction between the amplitude *A*_*n*_ of any of these linearly unstable modes and the amplitude *A*_1_ of the master mode corresponding to the eigenvalue *λ*_1_, yielding

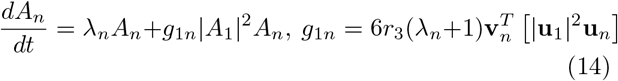

**FIG. 4.**
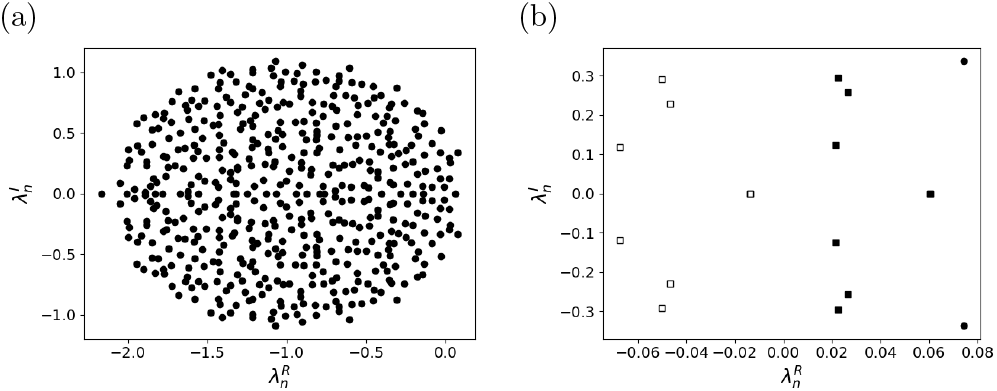
Learning in the presence of several unstable modes. (a) Spectrum of **L** for a network with *g* = 1.1 after learning *f* (*t*) = 2*f*_1_ cos *ωt* (*f*_1_ = 0.05, *ω* = *π/*10, *N* = 500). (b) Magnification of the unstable part of the spectrum. Nine modes have a positive real part (solid squares). Seven eigenvalues displaced by the interaction with the most unstable complex conjugate pair corresponding to 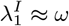 are also shown (empty squares) and all have negative real parts.

This interaction with the master mode keeps in check the seven modes *n* = 2, …, 8, by renormalizing their eigenvalues *λ*_*n*_ to *λ*_*n*_ + *g*_1*n*_ |*f*_1_| ^2^ which have negative real parts, as shown in Fig. 4b. The eigenvalue displacements depend on the function amplitude *f*_1_ and should be large enough to render negative the real parts of the *λ*_*n*_s. This prevents nonlinear restabilisation for small *f*_1_. Indeed, we found that learning fails for the matrix shown in Fig. 4 with *g* = 1.1 and *f*_1_ ≤ .05 as predicted by Eq. (14). This also agrees with previous results [7, 9] showing that forcing should be strong enough to suppress chaos, as intuitively expected.

The understanding of ESN that we have developed 5here, should hopefully help the design of actual ESN [2]. At the same time, it provides a theoretical framework to understand other learning approaches beyond ESN, including those that modify network connection strengths (**M**) in addition to output weights (**w**) [4, 5, 21]. We also note that networks with low-rank connectivity have recently been proposed to account for neural dynamics in several contexts such as discrimination tasks [22] or go-response task [23]. In effect, our analysis is complementary to these recent works in showing that feedback, in the regime we have considered, reduces the dynamics of a large random networks to that of few interacting modes, namely to the dynamics of a low-rank network. It also shows how to design low-rank networks with more general dynamics than those considered so far. It will be interesting to see whether real neural systems [24], such as the reciprocally interacting thalamus and cortex [18], have taken advantage of this design.

## END MATTER

### Analytic description of learning in linear networks

When the rate function is linear *r*(*x*), learning can be precisely understood, as we briefly show here, building upon previous works [16–18]. The problem is actually a particular case, for a linear feedback of rank 1, of the classical “pole placement” problem thoroughly studied in control theory [25, 26]. The forced solution **x**(*t*), needed to compute the correlation matrix and the r.h.s. of Eq. (3) obeys for a linear rate function *r*(*x*) = *x*,

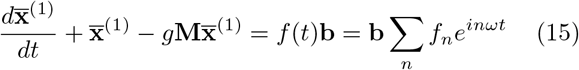

where we have introduced the Fourier decomposition of the function *f* to be learnt, with 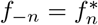 for *f* (*t*) to be real-valued. The forced solution is readily obtained as,

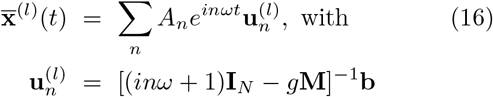

It provides an explicit expression of Eq. (3) for **w**,

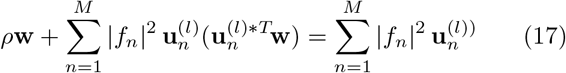

where denotes complex conjugation. The vectors 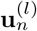 only span a subspace *U* of dimension *M* of the whole activity space of dimension *N > M* . Therefore, the correlation matrix is of rank *M* and **w** is underdetermined without regularization (*ρ* = 0). In order to more precisely see it, it is useful to imagine splitting **w** as **w** = **w**_*∥*_ + **w**_*⊥*_ in its parallel and perpendicular component to the sub-space *U*, as **w** = **w**_*∥*_ + **w**_*⊥*_ . Then, Eq. (17) reduces for the orthogonal component to *ρ***w**_*⊥*_ = 0. Thus, any finite regularization lifts the undeterminacy of **w** and imposes that it belongs to *U*, as also the r.h.s. of Eq. (17). This may be intuitively clear since the regularization imposes that the norm |**w**| = |**w**_*∥*_| + |**w**_*⊥*_| is minimal and that only **w**_*∥*_ contributes to solving Eq. (17). Since **w** = **w**_*∥*_ belongs to U, we can write it as a linear combination of the vector 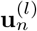,

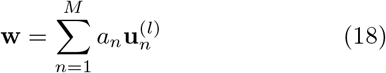

The *a*_*n*_ are the solution of the *M* × *M* system obtained by identifying the coefficients of each 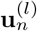 on the two sides of Eq. (17),

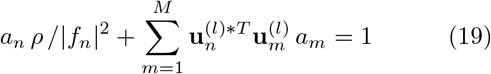

The very small regularization *ρ* in Eq. (19) has a negligible effect for a small number of modes *M* and the first term in Eq. (19) can be dropped. Eq. (19), then simply reduces to the conditions 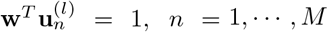 which implies that the 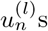 are eigenvectors of **L** (Eq. (4)). As a consequence, Eq. (16) for 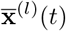 does not only provide a solution of the forced dynamics but also one of the autonomous linear dynamics. The projection of this linear solution by the readout **w** approximates the function *f* (*t*) when the amplitudes *A*_*n*_ in Eq. (16) are the amplitudes of the Fourier modes of *f* (*t*), *A*_*n*_ = *f*_*n*_.

### Explicit computation of normal forms

We detail here the computation of normal form equations and coefficients on the example of the single frequency sine function (Eq. (7)). As stated in the main text, after learning, **L** has two c.c. eigenvalues *λ*_1_ and 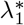 close to *iω* and −*iω*, with right eigenvectors **u**_1_ and 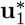 and left eigenvectors 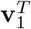 and 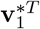,

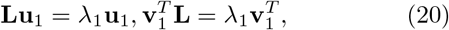

with the corresponding equations for 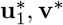 and 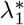. The eigenvectors obey the orthogonality relations 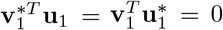 and we choose the normalization conditions 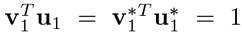. We search in perturbation for a small amplitude solution of Eq. (1) under the form **x** = [*A*_1_(*t*)*e*^*iωt*^**u**_1_ + *c*.*c*] + **x**_*p*_. The amplitude *A*_1_ evolves on a slow time scale controlled by the small real part of 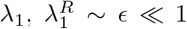, while |*A*_1_| ∼ *ϵ*^3*/*2^ and **x**_*p*_ ∼ *ϵ*^3*/*2^. Substitution into Eq. (1) gives for **x**_*p*_,s

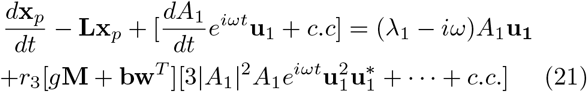

where the second term on the r.h.s. comes from the expansion of the rate function and the vector power is meant component-wise. We have only explicitely written the lowest-order “resonant” term proportional to exp(*iωt*). It gives rise to a secular term in **x**_*p*_ unless **x**_*p*_ has no component on **u**_1_ when written in the basis of the (right) eigenvectors of **L**. In other terms, it should be orthogonal to **v**_1_. Multiplying both sides of Eq. (21) on the left by 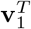 gives Eq. (5) with the expression (6) for the constant *g*_*A*_.

A similar computation provides the expressions of the coefficients *g*_*ij*_ for the normal form in the two-frequency case (Eq. (10)),

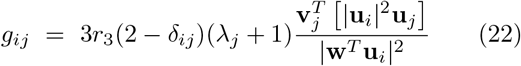

where the **u**s and **v**s are the right and left eigenvectors of **L** associated to the eigenvalues *λ*_1_ and *λ*_2_, as indicated by their subscript, They are normalized such that 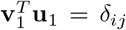, where here and in Eq. (22), we have used the Kronecker symbol, *δ*_*ij*_ = 1 if *i* = *j*, and 0 otherwise.

